# A study of Mutation in ATP7B gene and its correlation with clinical phenotype and radiological features in Wilson Disease patients

**DOI:** 10.1101/2021.01.21.427561

**Authors:** Jasodhara Chaudhuri, Samar Biswas, Goutam Gangopadhyay, Tamoghna Biswas, Jyotishka Datta, Atanu Biswas, Amlan K Datta, Adreesh Mukherjee, Atanu K Datta, Avijit Hazra

## Abstract

**Introduction:** Wilson Disease (WD) is an autosomal recessive disease caused by mutations in the ATP7B gene. Clinical manifestations of WD are variable. Identification of prevalent mutations in a given population is necessary to provide mutation-based molecular diagnosis. Previous studies have detected common mutations in this part of the world and our study aimed to correlate genotype with clinical and radiological features.

**Methods:** A descriptive cross-sectional observational study was conducted over a period of two years in a tertiary care hospital and neurology referral unit of Kolkata, India. All WD patients within the study period and meeting the inclusion criteria were included. Demographic data collection, clinical examination and relevant laboratory investigations were done. Magnetic resonance imaging of brain and cognitive assessment by Mini Mental Score Exam (MMSE) were also performed. Blood was collected for genetic analyses. PCR-Sanger sequencing of exons 2,4,6,8,10,14,16,18 of ATP7B gene was done based on previous reports of mutation hotspots of ATP7B gene for WD in Eastern India. Genotype phenotype correlation was attempted using two supervised machine learning methods, viz. logistic regression with an elastic-net penalty and the random forest.

**Results:** Of 52 WD patients were included in the study, 57.7% were males. The mean age at diagnosis was 13.96 years. Majority (61.8%) of the patients had dystonia on presentation, followed by dysarthria (41.2%), tremor (17.6%) and ataxia (11.8%). The mean MMSE and Frontal Assessment Battery score were 23.74 and 10.63 respectively and both were lower than the normal baseline values.Out of the total cohort of 52 patients,15(28.8%) harbored previously reported common mutations from this part of the country. Of the 15, 12 had the same mutation of c.813C>A(p.cys271Ter).The presence of common mutationswas associated with several distinct clinical phenotypes in the mathematical models but larger sample sizes are needed to corroborate the correlation.

**Conclusions:** WD patients in eastern India have significant genotypic and phenotypic diversity. Further studies with larger samples and screening of remaining exons are warranted.

## Introduction

Wilson disease (WD) caused by the mutation of P type ATPase gene (ATP7B) is an inherited inborn disorder of metabolism.Impaired ATP7B functioning leads to various neuropsychiatric manifestations of Wilson disease[1]. Currently, ATP7B is the only identified gene known to cause Wilson disease and mutations in this gene have been reported in almost all exons. Mutation screening has revealed a large number of defects covering almost the entire ATP7B, but efforts to correlate a specific mutation to a particular phenotype have been complicated by factors such as compound heterozygotes, large number of different mutations, limited number of patients and clinical heterogeneity even between affected siblings bearing the same set of ATP7B mutations[2]. The most consistent genotype–phenotype correlation in Wilson disease is that the most severe, early-onset disease with predominantly hepatic presentation is associated with mutations causing absent ATPase activity[3].The aim of the presentstudy was to study the mutations within ATP-BD of ATP7B gene in a regional Indian cohort with WD and to evaluate any potential correlation between genotype and phenotype.

## Materials and Methods

A descriptive cross-sectional observational study was conducted over a period of two years in a tertiary care hospital and neurology referral unit of Kolkata, India. The study was approved by the institutional ethics committee for human research and conformed to the principles of the Declaration of Helsinki. All patients gave informed consent prior to inclusion in the study. For children aged less than eighteen years, consent was obtained from their legal guardians. Subjects were enrolled from the ward and Movement Disorder clinic of Department of Neuromedicine, Bangur Institute of Neurosciences, Kolkata from January 2018 to December 2019.All WD patients aged five years and above attending our institute within the study period and meeting the inclusion criteria were included in the study. The diagnosis of WD was based on Sternlieb’s criteria [4] characterized by suggestive clinical features with evidence of positive Kayser–Fleischer (KF) ring, low serum ceruloplasmin and high urinary copper excretion. KF ring negative hepatic cases had no clinical or lab evidence of the etiology of other disease.Patients with only hepatic Wilson disease and those with preexisting neurological conditions were excluded.The clinical features were recorded in a pretested semistructured questionnaire.

The Neurological Involvement Score (NIS) was designed to capture the spectrum of neurological involvement of WD patients across cognition, behavior and motor domains. NIS has been devised following the Tier 2 of the Global Assessment Scale (GAS) for WD [5]. The development and validation of the NIS in the Indian population has been described previously [6]. Radiological data interpretation was done from magnetic resonance imaging (MRI) of brain after noting the anatomical distribution of the abnormalities.A grade was given for the severity of the change in signal intensity(0=Normal,1=mild,2=moderate,3=severe). The extent of disease was graded from 0 to 8 according to the area involved (a score of 1 was given for each of the following areas involved-lentiform nucleus,caudate nucleus,putamen,pons,midbrain,thalamus,cerebellum and atrophy or involvement of white matter)[7]. Cognitive assessment was done by Mini-Mental State Examination (MMSE) for adults and pediatric MMSE for children aged uptosixteen years.Frontal Assessment Battery (FAB) was used to assess the cognition also as the main cognitive dysfunction in Wilson is related to the subcortical cognitive networks [8].

In this study we tried to assess hotspot mutations by selecting few alleles which were known for mutation in our geographical region. We obtained the known mutations in our population by searching the Wilson Disease Mutation Database, the Human Gene Mutation Database, the UniProt (Universal Protein Resource) accession of the *ATP7B*-encoded protein, P35670, and the existing literature on mutant alleles found in East Asian population.

It has been previously suggested that exons 8, 12, 13, 15, 16, and 18 are hot spots for mutations in Indian WD patients[9]. The previous largest study from eastern India [6] identified exons 2,4,6,8,10,14,16,18 as common exons for mutations in WD patients of this region. Peripheral blood sample was collected from the included subjects and PCR-Sanger sequencing was done to screen these common exons for mutation.

### Detailed plans for statistical analysis

Differences in genotype frequency across the categorical variables, viz. disease manifestation, gender, consanguinity andKF-ringwas examined by chi-square test of homogeneity, while an analysis of variance (ANOVA) and t-test were used to testfor differences inage at diagnosis, serum ceruloplasmin across different groups. A significance threshold of p-value of less than 0.05 was used to find non-null associations and group-differences.Futhermore, the strength of associationbetween lesion count on MRI and FAB score was explored by constructing scatter plot and calculating Spearman’s rank correlation coefficient *ρ*, that measures the degree of monotone relationship between two variables.

The Kolmogorov-Smirnov goodness-of-fit test, used to test the assumption of Normality of the numerical variables, revealed that Age, AgeAtDx, CERU did not show any for any significant deviation fromNormality while MMSE, MRI_HI, MRI_LOC and FABScoreappeared to have higher skewness than a fitted Normal distribution. For a deeper insight on the relationship between the predictor variables with presence or absence of common mutations, we utilized two popular supervised machine learning methods, viz. logistic regression with an elastic-net penalty [10] and the random forest [11]. We discuss the methods in greater details in our Results section.

### Statistical softwares used

- Statistica version 6 [Tulsa, Oklahoma: StatSoft Inc., 2001]
- MedCalc version 11.6 [Mariakerke, Belgium: MedCalc Software 2011]
- R Core Team (2019). R: A language and environment for statistical computing,Vienna,Austria.

## Results

In this study, we report the findings based on 52 subjects used for analysis and interpretation of results. Among these 52 patients, 30 (57.7%)were male and 22 (42.3%) female.The age distribution of the study population ranged between 10 years and 38 years with the mean being 19.83 years. The mean age at diagnosis was 13.96 years. Majority (61.8%) of the patients had dystonia on presentation, followed by dysarthria (41.2%), tremor (17.6%) and ataxia (11.8%).

For ease of comparison, we consider a further sub-division:the first, larger group was the whole population or the cohort under consideration and the second sub-group, subsumed within the first group,consisting of individuals possessingcommon mutation.By doing so, we tried to compare whether the group with common mutation genotype was having different correlations with the demographic variables. Out of the total cohort of 52 there were 15(28.8%) individualswith common mutations.Table 1 shows the descriptive statistics for the numerical covariates for the 15 individuals with common mutation.

**Table 1:** Descriptive Statistics for Numerical Variables – Cohort with Common Mutations [N = 15].

| Variable | N | Mean | Median | Minimum | Maximum | Upper Quartile | Lower Quartile | Std. Dev | Std. Error |
| --- | --- | --- | --- | --- | --- | --- | --- | --- | --- |
| Age | 15 | 19.2 | 17 | 10 | 32 | 14 | 24 | 6.394 | 1.651 |
| AgeAtD<sub>x</sub> | 1<br>5 | **13** | 12 | 7 | 24 | 10 | 16 | 4.884 | 1.261 |
| MMSE | 1<br>4 | **22.7**<br>**9** | 24 | 12 | 28 | 23 | 26 | 5.132 | 1.372 |
| CERU | 1<br>3 | **12.2**<br>**4** | 11 | 7.1 | 23 | 8 | 16 | 4.876 | 1.352 |
| MRI_HI | 1<br>5 | **1.53** | 1 | 1 | 3 | 1 | 2 | 0.743 | 0.192 |
| MRI_L<br>OC | 1<br>5 | **3.27** | 3 | 2 | 6 | 2 | 4 | 1.28 | 0.33 |
| FABScore | 1<br>5 | **11** | 12 | 8 | 16 | 9 | 12 | 2.507 | 0.647 |

At a genotypic level, of these 15 subjects with common variants,12 had mutation of c.813C>A(p.cys271Ter).The novel mutations areNM_00053.3 C.3389C>7 (uncertain),C.3389C>T, C.3722C>T and C.3389C>C/T.The clinical finding that was associated with two new mutations was Tics. However, more sample size is required to statistically analyze the clinical variation of these new mutations. The mean MMSE that is the screening cognitive score in the total cohort was found to be 23.74 and the mean Frontal Assessment Battery score in the entire cohort was found to be 10.63 and both were lower than the normal baseline values.The mean ceruloplasmin value was 12.24, lower than normal, MRI hyperintensity score mean value is 1.67 and the location score is 3.39.

In our study, we also found that lower ceruloplasmin values are associated with younger age group. This is visible in the correlation plot showing the association between variables and the higher numerical values are associated with higher chance of correlation. So according to this figure FAB score and MMSE shows a higher correlation value. Ceruloplasmin and FAB score shows the lowest correlation value. Rest have intermediate correlation values.[Figure1]

**Figure 1.**
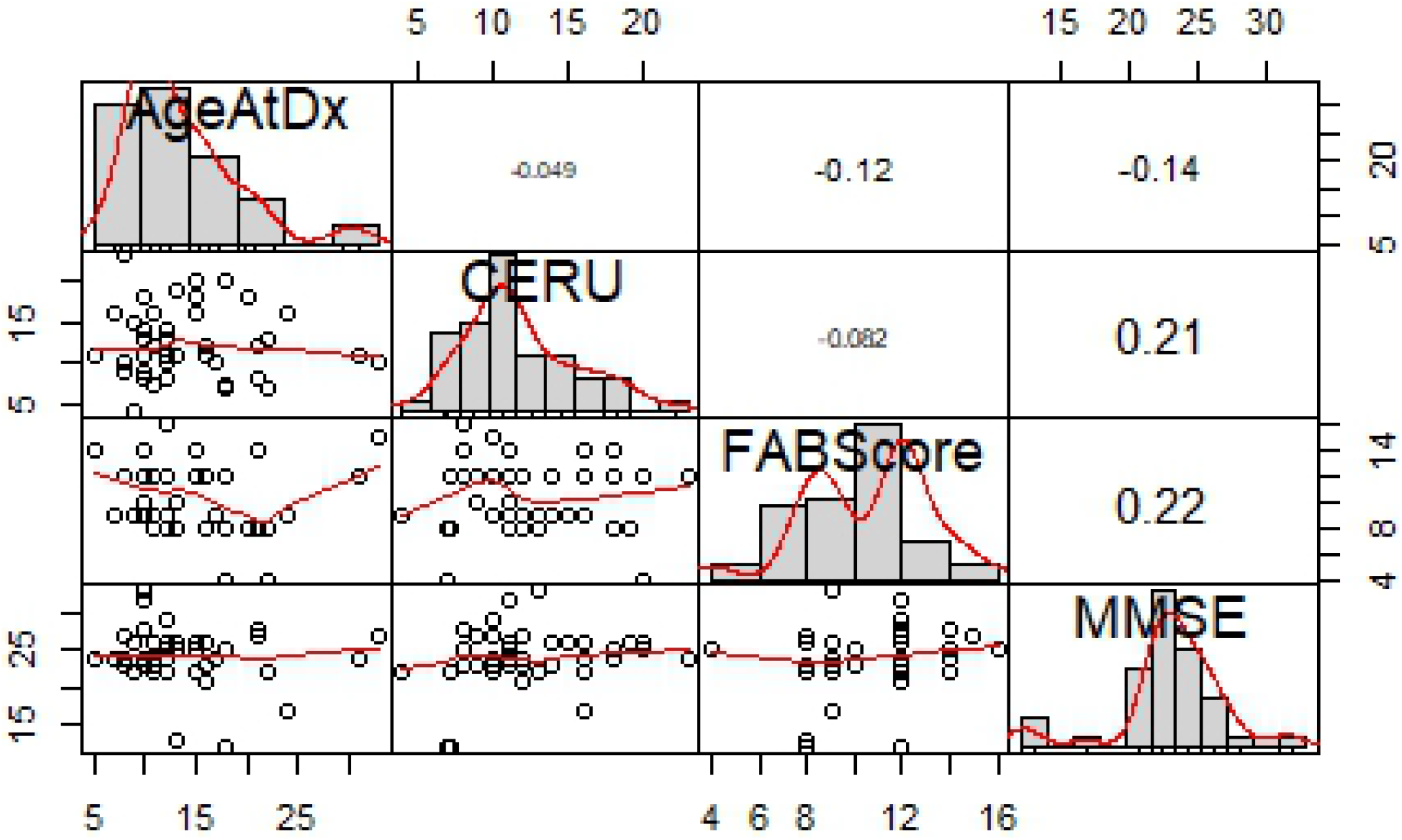
Histogram and scatter plots for numerical covariates.

Next, we show two figures showing association between factors using by the Chi-square p-value as well as the Cramer’s V measure of association. We split the association matrix into two halves for the sake of presentation, with ‘common mutation’ being the common variable on both the plots. The first figure suggests strong association between different clinical variables but not so much with Common Mutation, and the second figure suggests a moderateassociation of Common Mutation with Parkinsonism, Dystonia, Depression and Gait.[Figures 2 and3]

**Figure 2.**
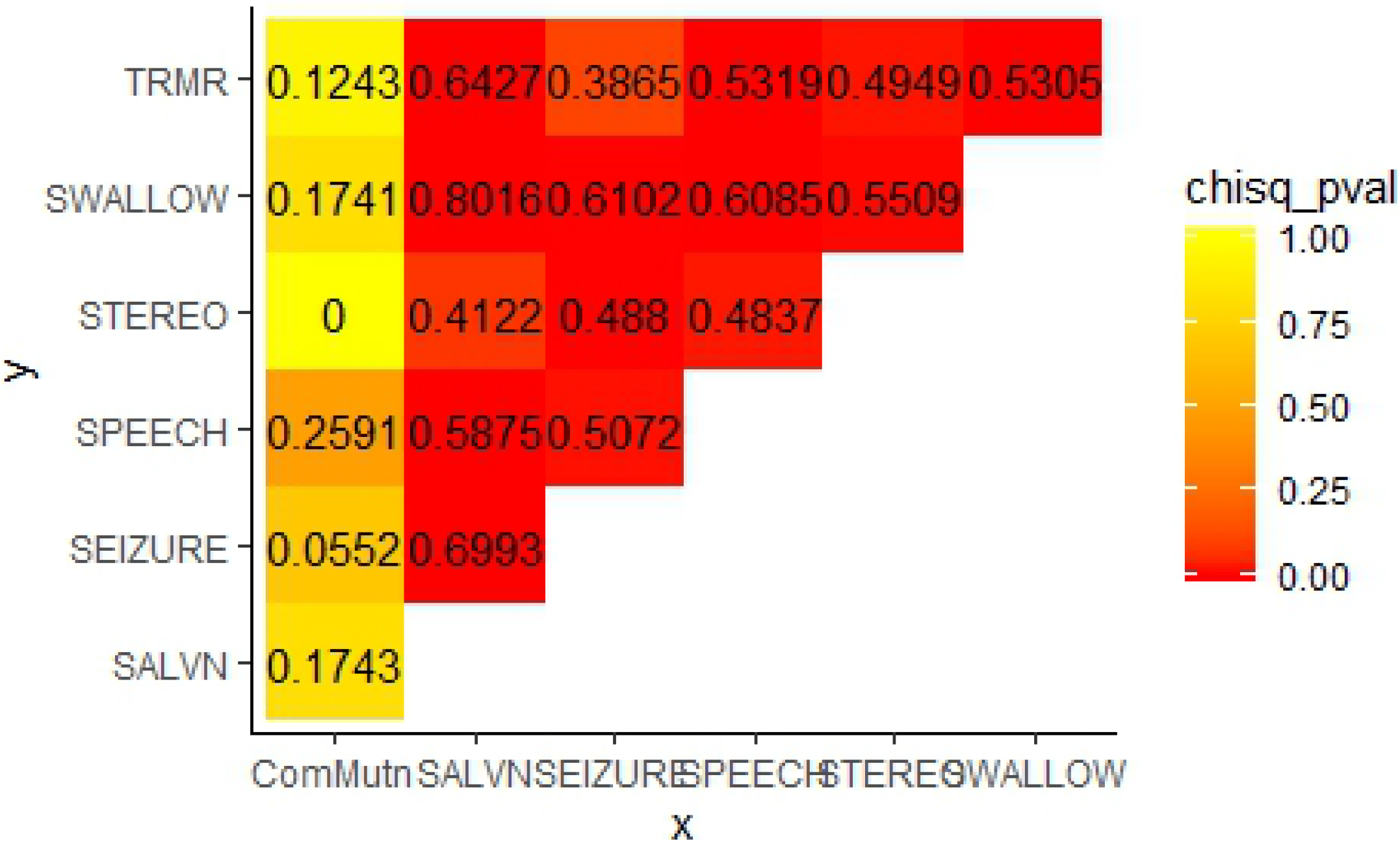
Association measures and Cramer’s V - part 1.

**Figure 3.**
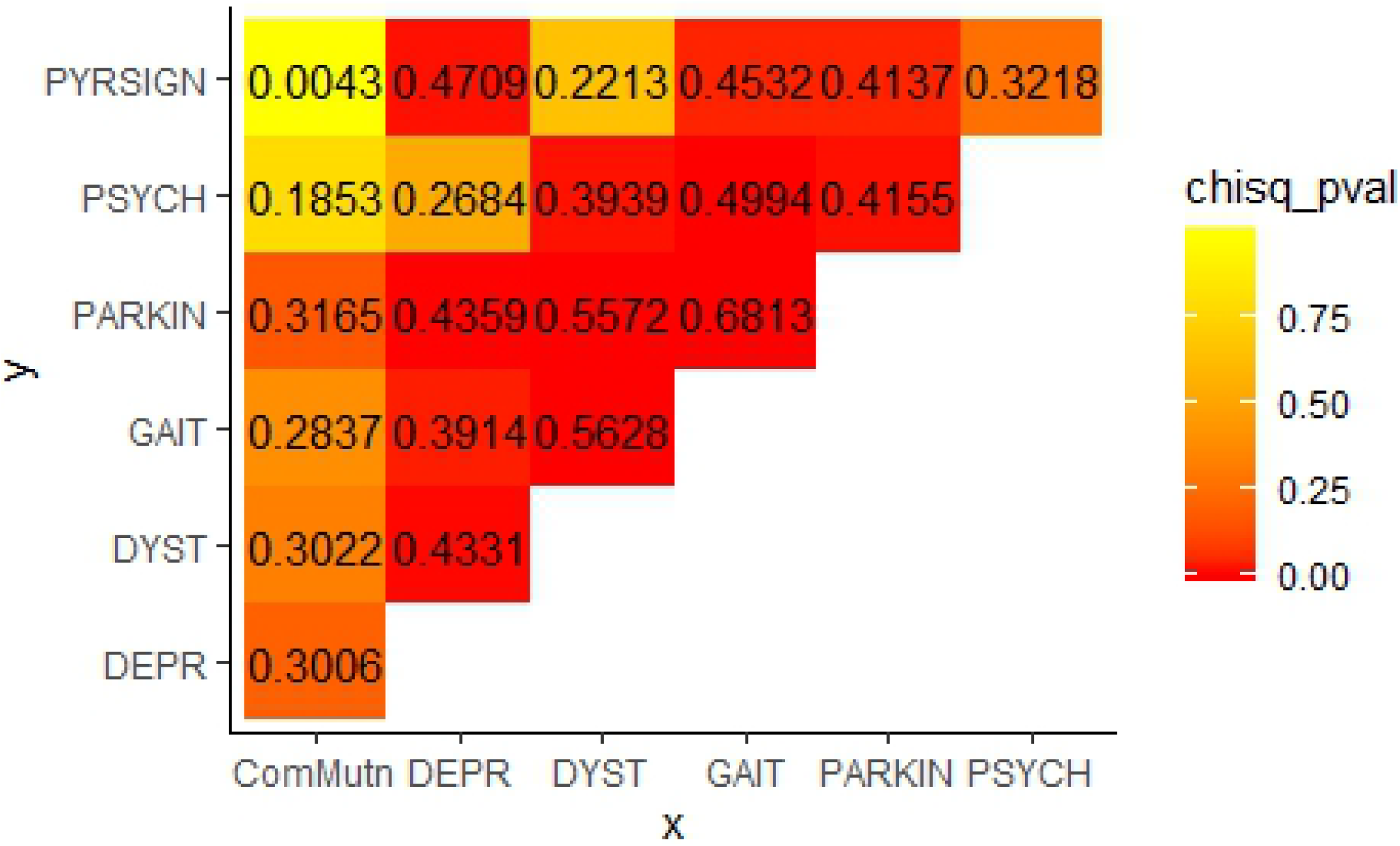
Association measures and Cramer’s V - part 2.

### Comparison across subgroups

Figures 4 and 5 belowshow how the four numerical variables *viz*. Age at diagnosis, Ceruloplasmin, MMSE (cognitive scale), and FAB scores vary across the groups – with or without common mutations. It seems from these boxplots that the differences for these variables between different levels of Common mutation are not significant, indicated by the substantial overlap of the probability regions.The boxplots suggest no significant difference in age distribution or ceruloplasmin levels exists between the two groups(1 depicts common mutation group and 0 depicts others).[Figures 4 and 5]

**Figure 4.**
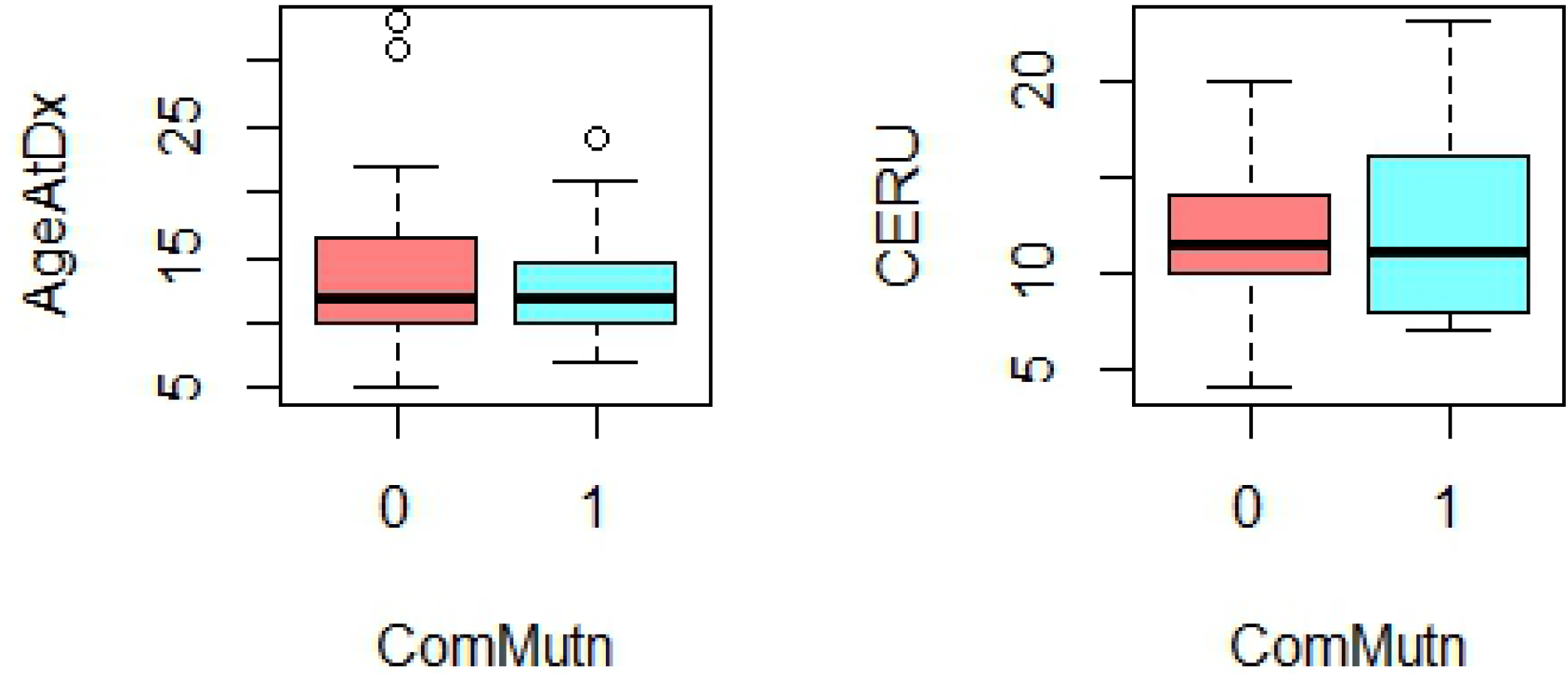
Boxplot showing AgeatDX and CERU between groups.

**Figure 5.**
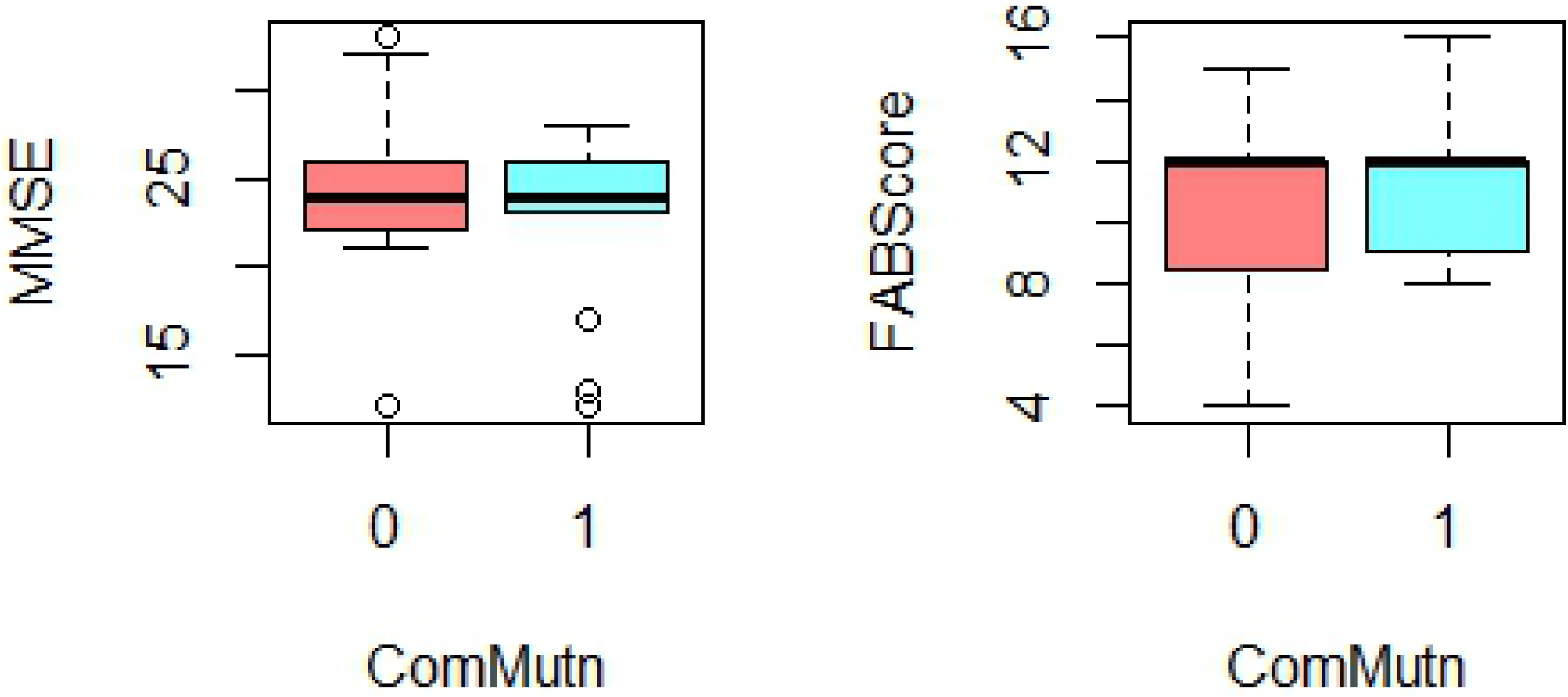
Boxplot showing MMSE and FABScore between groups.

### Predicting Class Labels using Supervised Learning

Next, we use a supervised learning approach to evaluate the performance of two popular statistical machine learning methods: (a) the penalized logistic regression and (b) the random forest to predict common mutation. The key idea behind supervised learning algorithm is to divide the data into two parts - training samples and test samples, and fit the model on one training samplesand evaluate its performance such as prediction or inference on the remaining part, the test

data.Supervised learning method prevents overfitting –fitting the training data too closely to generalize to do well on new observations [12]. In our context, this means we estimate the parameters of the random forest or logistic regression by minimizing the error from the training samples via cross-validation and the performance is reported on the test data. For the results reported in this paper, we use a 50-50 split, resulting in 25 observations for training, 26 observations for the test data.

First, we briefly describe the two supervised learning approaches used in this paper:

### Penalized Logistic Regression

We fit a logistic regression for explaining common mutation as a function of the available predictors. A logistic regression fits the log-odds of the probabilities as a linear function of the predictors, i.e.

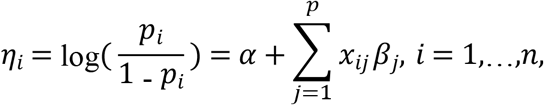

where the *x*_*ij*_ is the i-th observation for the j-th predictor.The usual regression approach would run into a problem here as we have a low-rank data, i.e. the number of predictors is more than the number of columns in our data frame, as each ordinal factor has to be recoded into a baseline category, and one predictor for each different level. To solve this, we use a penalized regression called the elastic net [10, 12, 13]. The elastic net regression minimizes the penalized log-likelihood:

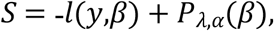

where *l*(*y,β*) is the log-likelihood, given below, and *P*_*λ,α*_(*β*) is the penaltyterm:

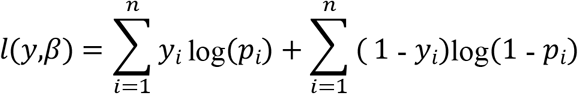

and the elastic net penalty term is given by:

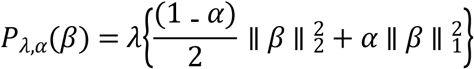

We use the same R package ‘glmnet’ for implementing the penalized logistic regression.[Figure 6]

**Figure 6:**
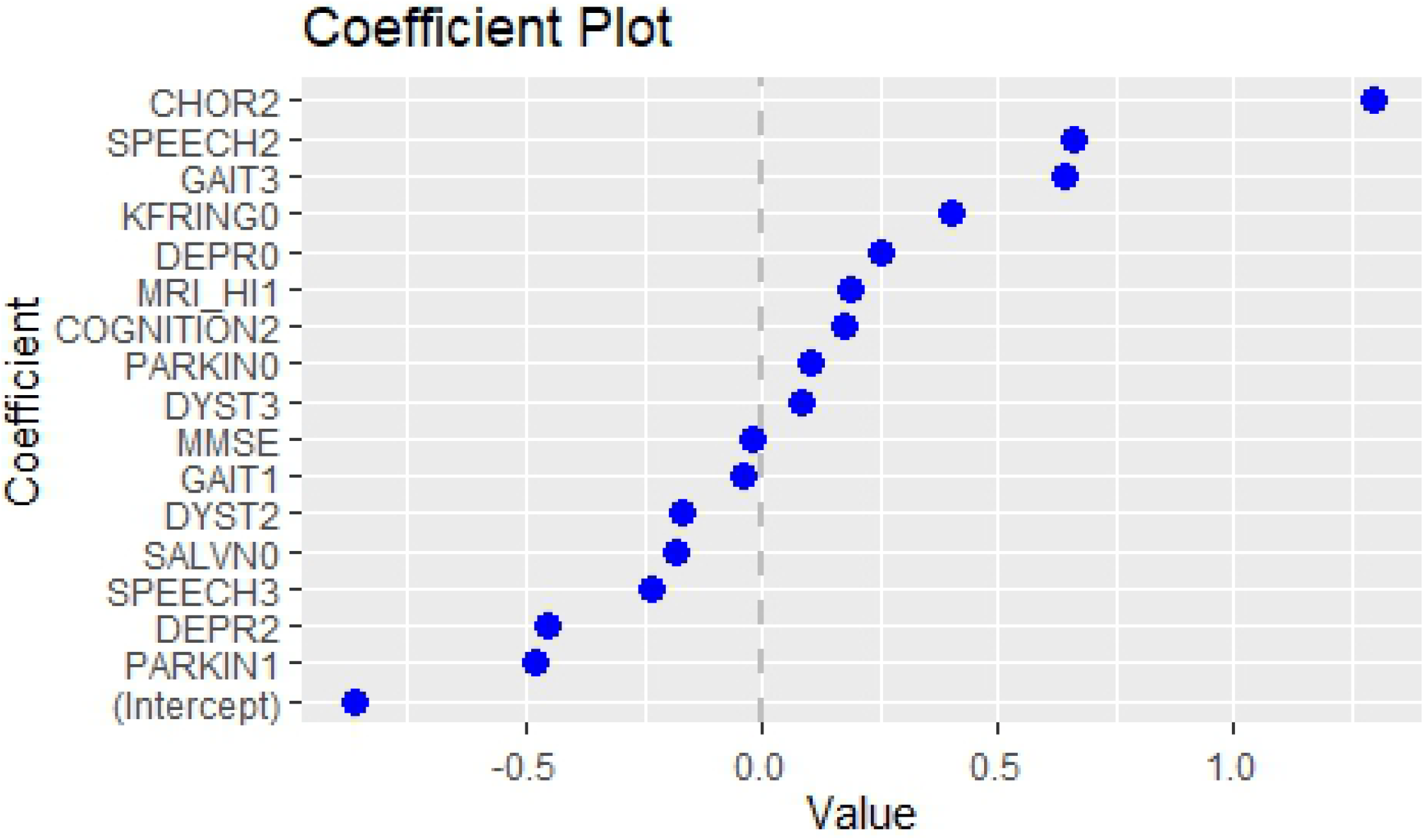
Coeffeicient Plot

### Random Forest

The random forest [12,14,15], on the other hand, is an ensemble method where many classification trees are built on resamples of the training data that are then averagedfor obtaining predicted class labels. For a single regression tree method, a large decision tree is grown depending on recursivepartitions of the predictor space that yields a small residual sum of squares on training data. The random forest is fitted by minimizing an appropriate error metric on training samples with an added penalty term that prevents over-fitting by controlling the size of the tree. Being an ensemble of many space-partitioning classificationtrees, random forest tends to outperform linear methods like penalized logistic regression when the true relationships are non-linear in nature.[Figure7]

**Figure 7.**
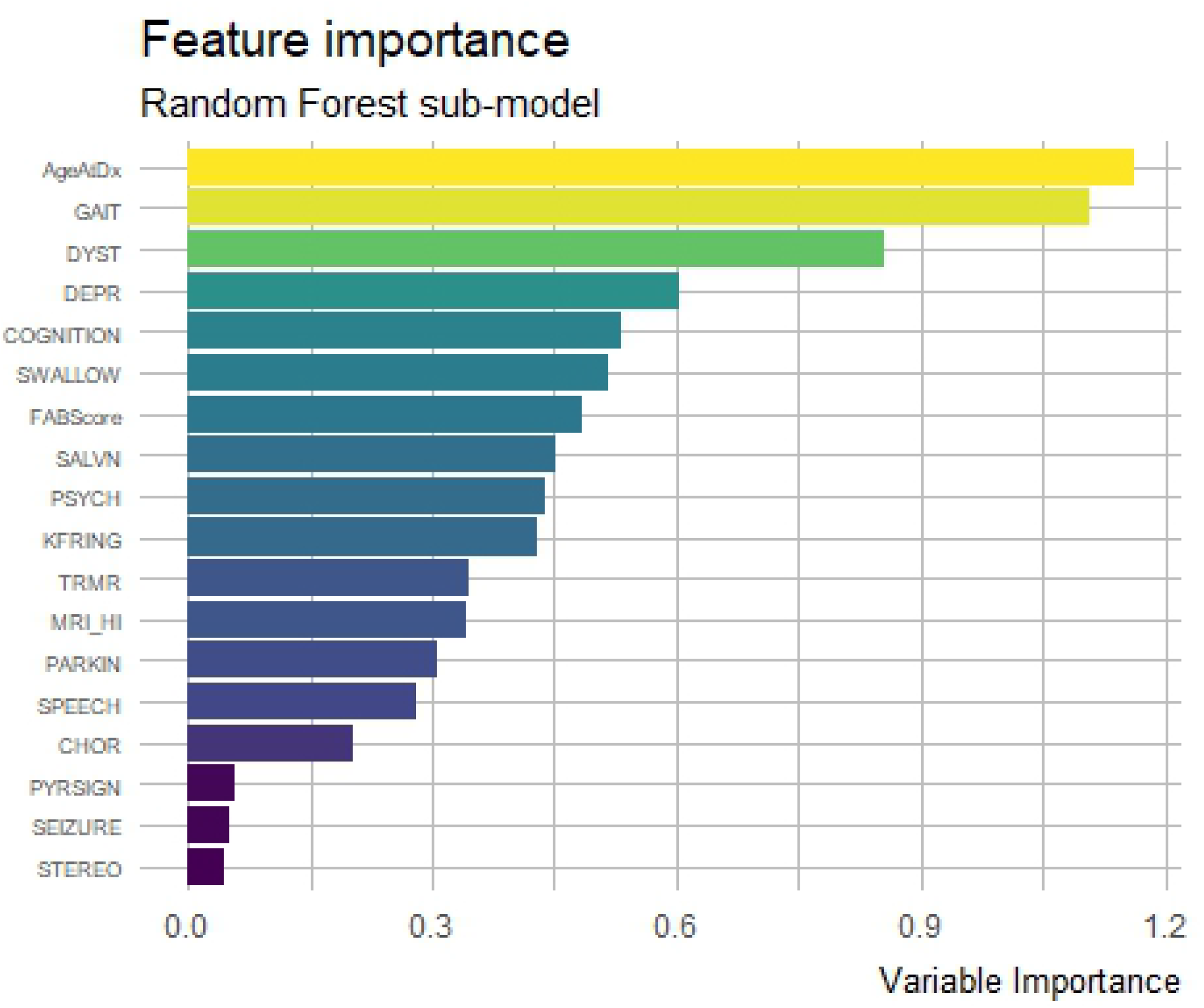
Feature importance for random forest classifier

### Feature Importance: Random Forest

Feature importance or variable importance for random forest is a measure of the degree of contribution by each predictor to the classification task. Figure 9 visualizes ‘variable importance’ for the random forest, showing which features make the greatest contribution in predicting common mutation. For ourrandom forest classifier, Gait and Age at Diagnosis has the greatest contribution, followed by Dystonia and Depression. The order of importance is not the same as that from the logistic regression model, which is expected since the random forest fits a space-partitioning model instead of a linear model.

Now, we show the relative performance of the two methods: logistic regression and random forest. The first two tables show the predicted class labels versus the true class labels. For our training-test random split, the probability of correct classification for Random Forest was 0.576923, compared to the misclassification probability of logistic regression being 0.538462.The cross-tabulated data for true and predicted class-labels are given below, with ‘0’ and ‘1’ denoting absence and presence of common mutation on an individual. These tables show that while the classification algorithms are good at predicting the absence of common mutation (with true positive rates being 0.7647 and 0.8823 for logistic classifier and random forest), but poorly perform in predicting the ‘1’ labels or presence of common mutations (see Tables below).

| Logistic regression classifier |  |  |
| --- | --- | --- |
|  | True Class Labels Test Data |  |
| Predicted Class Labels | 0 | 1 |
| 0 | 13 | 8 |
| 1 | 4 | 1 |

| Random forest classifier |  |  |
| --- | --- | --- |
|  | True Class Labels Test Data |  |
| Predicted Class Labels | 0 | 1 |
| 0 | 15 | 9 |
| 1 | 2 | 0 |

## Discussion

The present study included 52 patients with neurologic manifestations of Wilson disease. The age distribution in our patient population ranged from 10 yearsto 38 years with the mean age being 19.67 years. The mean age at diagnosis in the study population was 13.96 years. In their study on late onset Wilson disease Ferenci et al [16] showed majority of patients had onset below 30 years of age and this is corroborating with our study. Out of the total cohort of 52 patients,15(28.8%) harbored previously reported common mutations from this part of the country. Of the 15, 12 had the same mutation of c.813C>A(p.cys271Ter). The biggest study from Eastern India by Mukherjee et al[6] in 174 patients reported about 58 different coding mutations of which 21 were novel and 37 previously reported variants with common mutations accounting for 24%of the total mutations in Eastern India. However, in our study we did not do the promoter reporter assay to determine whether the genetic variants in the promoter and 5’UTR could affect WD gene expression.A study by Gupta et al[17] in 2005 found out that c.813C>A (Cys271Stop) and c.1708-1G>C were found to be associated with 23 and 12 WD chromosomes, respectively amongst a total of 64 patients. In 2007 the same study group Gupta et al. [18] published a study with 114 patients and found out that in addition to the previous mutations 8 new mutations were found in the population. Among the prevalent mutations Cys271Stop, which accounts for around 19% of the total mutations, has been detected earlier in the European and Turkish population, but not as a prevalent one. From their study it was concluded that for studying the mutations in the Indian population it would be important to screen exon 2 first, where we observed largest number of nucleotide variants including three prevalent mutations. Other large studies have also attempted to identify mutations in the north [9] and south [19] Indian populations. Screening for these prevalent mutations can be the first step of genotyping for carrier and presymptomatic sibling screening in populations from the three zones..In our study the commonest mutation found was c.813C > A mutation which is a non sense mutation found in Exon 2 and is also the commonest mutation in the eastern part of India. The previous studies failed to have definite correlation between the genotype and phenotype, and this could be attributed to presence of modifier loci in WD phenotype.

In our study we made two subcohorts –one group with common mutations and another one without common mutations.In the group with common mutations the MMSE and mean FAB score were 22.74 and 11 respectively. The group without common mutationshad a mean MMSE score of 24.67 and a mean FAB score of 10.5. There was no significant difference in the cognition between the two groups.FAB score and MMSE showed correlation amongst one another in our study but there was no correlation with genotype. There was no correlation of FAB score and MMSE with ceruloplasmin as shown in other studies. A higher lesion count on MRI was associated with a lower FAB score in all patients implicating that higher lesion load in MRI brain leads to lower FAB score and poorer cognitive function. The scatter plot of age versus ceruloplasmin suggested that a lower age of presentation was associated with lower ceruloplasmin values. This is in concordance with the study by Cheng et al[20], who showed that the lowest ceruloplasmin levels were found in the age group of 7-12 years. However, the ceruloplasmin levels were not found to be related to common mutation in our study. The presence of common mutations was more positively correlated to poor speech,dystonia,parkinsonism and depression more than the other predictor variable.The earlier study from Eastern India by Mukherjee et al. [6] concluded that the phenotypicspectrum emerging from a G2P matrix of Wilson disease was quite diverse, critical examination of the entire profile revealed that a few patients depicted significant differences in terms of their clinical presentation as compared to other patients in the same mutation cluster leading the authors to postulate that genetic alterations in other copper metabolism pathway genes or environmental factors might be responsible for phenotypic variability observed between patients who harbor identical/similar mutations in ATP7B.

The next aspect of our study was neuroimaging which was done by 3T MRI and there was no significant difference in the MRI scores both in terms of location of abnormalities and intensities in the common mutation group and the other group.The study on cranial imaging in Wilson by King et al. [7] showed that basal ganglia specially the putamen is the most affected part in Wilson disease followed by pons, medulla and cerebellum. In our study we tried to correlate the MRI lesion load with FAB score and found that a higher MRI lesion correlated with a lower FAB score that is a poorer cognitive outcome probably by affecting the frontal subcortical circuits.

Due to paucity of number of subjects we attempted to construct mathematical models to predict the relationship of genotype with phenotype that will be applicable to possible future studieswith larger populations.The goodness of fit model was done and showed that common mutation can have some correlation with phenotypic characteristics like chorea,speech and KF ring.Lastly, a classification performance model was done using elastic net penalized regression and random forestas supervised machine learning approaches. The probabilities of correct classification by our methods are 0.54 and 0.58 for logistic classifier and random forest, respectively, which suggests the predictive power of this model of common mutation correlating with phenotypic characteristics is marginally higher than by random chance alone.The Random Forest model also showed that the predictive value of some predictor variables are more associated with common mutation: these were younger age,gait,dystonia and depression. Overall, the performances of the two models suggest that it is possible to do better prediction with common mutations with a larger sample size, and more statistical power. To the best of our knowledge, ours is the first study from eastern India to explore mathematical models as a way to predict genotype phenotype correlation in Wilson disease patients. Further research with larger sample sizes should be conducted to better understand and corroborate our findings.

## Acknowledgement

Dr Paramita Bhattacharya, Genetic Laboratory Scientist, National Institute of Biomedical Genomics,Kolkata.

## Appendix 1

### A.1. Predicted class probabilities for penalized logistic classifier

The first panel on the figure shows the predicted probabilities and the actual prediction and the second shows the true labels vs the prediction.

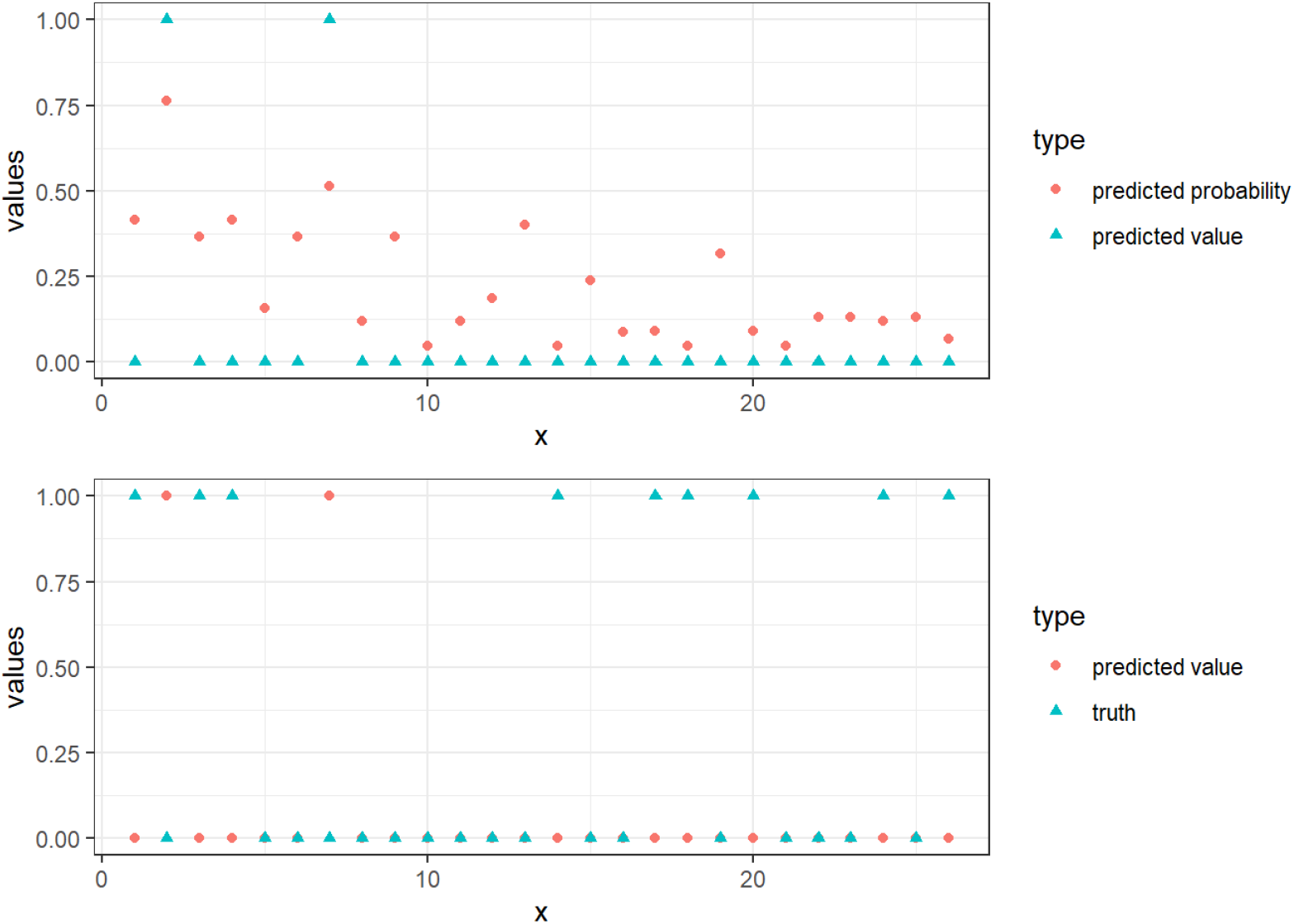

## Appendix 2

